# Discover the Maze-like Pathway for Glabridin Biosynthesis

**DOI:** 10.1101/2025.05.06.652391

**Authors:** Zhen Zhang, Wenqiang Li, Fanze Meng, Yanming Jin, Wentao Sun, Qina Kuang, Shichao Ren, Wei Liu, Liang Zhang, Lei Qin, Bo Lv, Haiyang Jia, Chun Li

## Abstract

Tailoring enzymes diversify plant metabolite scaffolds through complex, context-dependent modifications, generating maze-like biosynthetic networks that complicate metabolic pathway reconstruction. Here, we systematically deconstructed and reconstructed the biosynthetic network of glabridin, a valuable skin-whitening isoflavone from *Glycyrrhiza*. By integrating metabolic pathway mining with genome and 183 transcriptomes, we identified four functional routes among sixteen theoretical possibilities and uncovered a previously uncharacterized, ladder-like multi-route tailoring network. Reconstruction of such architecture in yeast revealed metabolic redundancy and interconnectivity confer unexpected robustness, enabling higher production efficiencies compared to a single-route design. Further modular engineering enabled the *de novo* biosynthesis of glabridin in yeast. Our work establishes a framework for reconstructing metabolic mazes and demonstrates that multi-route architectures can be harnessed to enhance the robustness and productivity of cell factories.

## Main Text

Tailoring enzymes(*1–3*) are essential in plant secondary metabolism, where they steer the synthesis of complex bioactive compounds by modifying a foundational metabolite scaffold. Unlike the initial assembly of a basic molecular framework, tailoring introduces a series of chemical modifications that determine activity, solubility, stability, and subcellular localization(*4, 5*). This post-assembly process is not only critical for generating the diverse array of natural products, but it also adds an additional layer of regulation that ultimately shapes the biological activity of these compounds. However, this tailoring complexity presents a challenge when attempting to reconstruct or engineer the biosynthesis of specific target compounds. The intricate sequence of reactions due to the promiscuity often gives rise to a metabolic network that resembles a maze—each step interwoven and highly context-dependent(*6, 7*). Such complexity makes it exceptionally difficult to fully map the biosynthetic pathways of certain specialized metabolites, especially those with multiple modifiable sites and obscure tailoring logic.

In the realm of flavonoid biosynthesis, more than 9,000 distinct compounds have been tailored from common precursors, representing one of the most structurally diverse classes of natural products(*8*). Among them, glabridin(*9*), a naturally occurring prenylated isoflavone, is renowned for its skin-whitening, anti-inflammatory, and anti-aging properties(*9, 10*). It commands a multimillion-dollar global market in high-end whitening cosmetics. However, glabridin is difficult to chemically synthesize due to the challenge of stereoselective construction of the C-3 chiral center and the chemoselective reduction of conjugated systems(*11*). Currently, the world’s supply chain for glabridin relies on low-yielding extraction and purification from the plant root of rare plant *Glycyrrhiza glabra L*, which only accounting for 0.08-0.35% of the roots’ dry weight(*10*). Microbial or synthetic biosynthesis of glabridin remains challenging due to a complex maze of post-tailoring modifications, and even its native biosynthetic pathway has not yet been fully elucidated.

In this study, we applied a metabolic pathway AI search algorithm integrated with experimental metabolic data to comprehensively elucidate and map a maze-like biosynthetic network of glabridin. By systematically mining the transcriptome and genome data of *Glycyrrhiza* species, we identified all key tailoring enzymes involved in glabridin biosynthesis. These enzymes were then used to reconstruct a multi-route synthetic pathway, in which a single set of promiscuous enzymes supports multiple parallel biosynthetic routes. Through this reconstruction, we uncovered distinct biosynthetic modes underlying glabridin formation and ultimately achieved *de novo* biosynthesis of glabridin in *Saccharomyces cerevisiae*.

### Glabridin pathway design and core nodes identification

To discover the biosynthesis pathway of glabridin, we employed a metabolic pathway search algorithm based on known enzymatic reactions in combination with a retrosynthetic algorithm derived from reaction rules(*12–14*), which enables pathway design by deconstructing target molecule and medicarpin (**8**) as key intermediates into simpler precursors in yeast step-by-step. Using this approach, we designed 16 potential biosynthetic pathways leading to the production of glabridin, involving L-phenylalanine (**1**) as precursor and 31 intermediate metabolites (Fig. 1a). Among these, 12 compounds are located at pathway branch nodes, and over 64% of the predicted pathways include more than six branch nodes. The presence of such diverse and numerous branch points significantly increases the complexity of glabridin biosynthesis.

**Figure 1.**
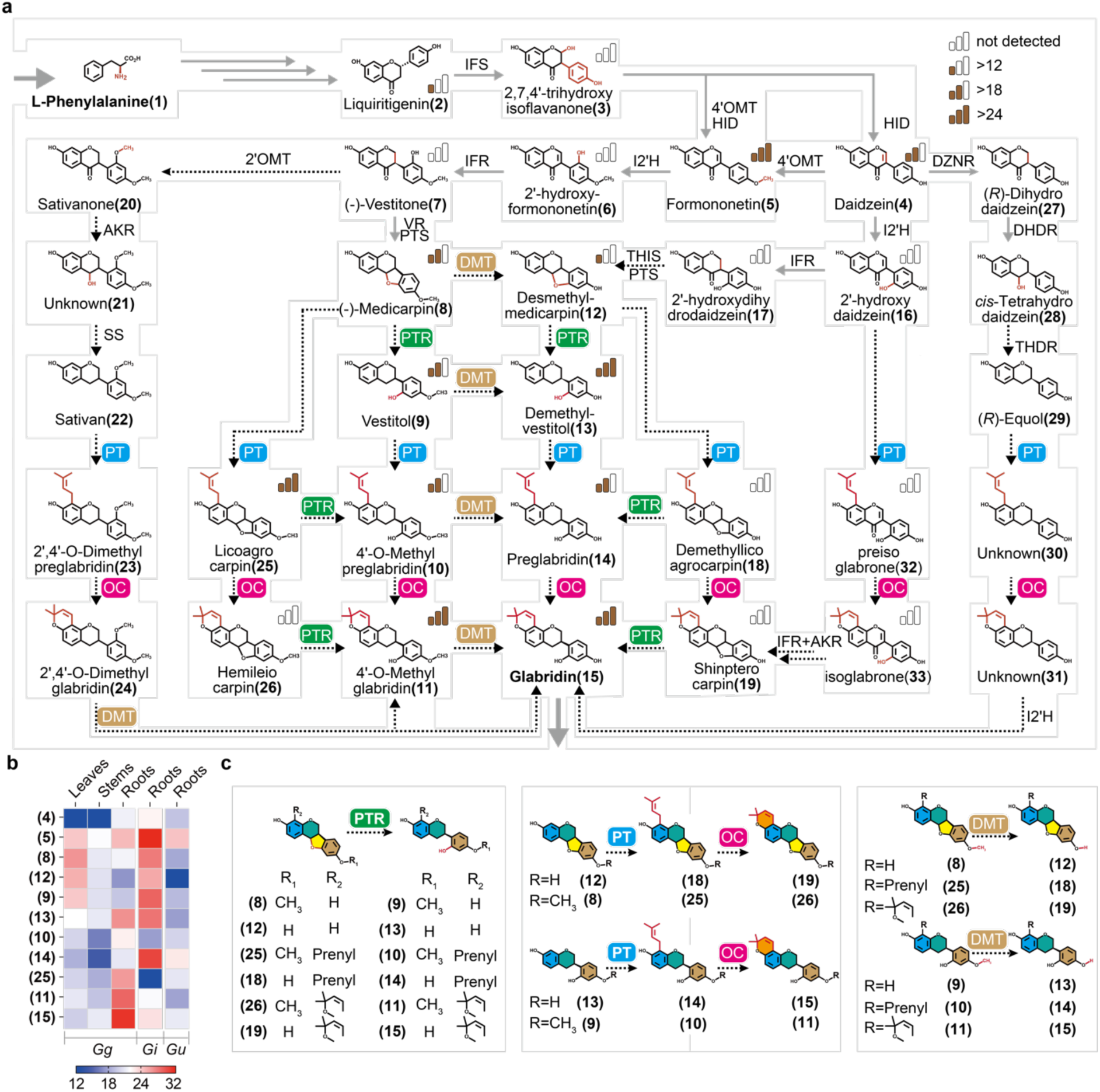
The maze-like glabridin biosynthesis pathway. a) Summary of predicted glabridin biosynthesis. b) Metabolomic profiling of different organs and species of *Glycyrrhiza,* including *Glycyrrhiza glabra* (*Gg*), *Glycyrrhiza uralensis* (*Gu*), and *Glycyrrhiza inflata* (*Gi*). c) The key tailoring steps in glabridin biosynthesis. Dash arrows in a. and c. represent uncharacterized step, gray arrows represent characterized steps, and the overlapping arrows represent a multi-step reaction. The color bar in b. indicates the log_2_-transformed peak area. Metabolite abundance is categorized into four levels based on accumulation in *G. glabra* roots: high (>24), medium (>18), low (>12) and undected. The bar chart in the upper right corner of each compound in a. indicates its dection in *G. glabra* roots, where the number of solid brown bars corresponds to the abundance level, and hollow bars indicate that the compound was not detected. Abbreviations not already defined in the text are as follows: isoflavanone 2’-O-methyltransferase (2’OMT), aldehyde ketone reductase (AKR), sativan synthase (SS), isoflavone synthase (IFS), 2,7,4’-trihydroxyisoflavanone 4’-O-methyltransferase (4’OMT), 2-hydroxyisoflavanone dehydratase (HID), isoflavone 2’-hydroxylase (I2’H), isoflavone reductase (IFR), vestitone reductase (VR), pterocarpan synthase (PTS), 2’-hydroxydihydrodaidzein synthase (THIS), daidzein reductase (DZNR), dihydrodaidzein reductase (DHDR) and tetrahydrodaidzein reductase (THDR).

To evaluate the most feasible biosynthetic routes among those predicted, we conducted metabolomic profiling of three *Glycyrrhiza* species: *Glycyrrhiza glabra*, *Glycyrrhiza uralensis*, and *Glycyrrhiza inflata* (Fig. 1a-b). Metabolomic analysis revealed the major metabolites, such as compounds **5**, **11**, **13**, **15** and **25**, are predominantly accumulated in the roots compared to leaves and stems. Glabridin was detected only in *G. glabra* and *G. inflata*, with the highest accumulation observed in *G. glabra*. Furthermore, several predicted intermediates (**4**, **8**, **9**, **10**, **12** and **14**) were also identified in *G. glabra*, providing support for their roles as key metabolic network nodes(*15*). Thus, through node-centric analysis of the metabolomic data, the predicted routs major incorporated with key intermediates, such as medicarpin (**8**), were further considered. Even though, 12 possible alternative biosynthetic pathways for glabridin still persist due to the diverse nature of tailoring reactions. This complexity complicates pathway design and hinders efficient metabolic engineering of glabridin.

To address this complexity, we simplified the pathway by categorizing and deconstructing glabridin biosynthesis into four classes of tailoring reactions: pterocarpin reduction, prenylation, prenyl cyclization, and demethylation (Fig. 1c). These transformations are likely mediated by four classes of tailoring enzymes: pterocarpin reductases (PTRs), prenyltransferases (PTs), oxidative cyclases (OCs) and demethylases (DMTs). Among these, PTRs, along with VRs and IFRs, belong to the class of NADPH-dependent reductases(*16–18*), sharing similar catalytic mechanisms and often exhibiting high sequence similarity(*18*), which poses a challenge for accurate PTRs identification. PTs and OCs enzymes were found to broaden the range of possible intermediate metabolites. In addition, DMTs catalyze critical branch-forming reactions, with the high branching complexity, which direct the metabolic flux toward different outcomes. The four tailoring reactions can proceed in different sequential orders through various branching nodes, all ultimately leading to produce glabridin (Fig. 1a).

### Systematic omics analysis for enzyme discovery

To identify suites of candidate genes from *G. glabra* that can be used to reconstitute glabridin biosynthesis, especially for the four classes of tailor enzymes, we used systematic seasonal and organ specific transcriptome, genome mining and co-expression analysis (Fig. 2). We first collected wild perennial samples of *G. glabra* from natural habitats in northwest of China and performed genome sequencing and assembly (Fig. 2a). We assembled a chromosome-level *G. glabra* genome with a total size of 415 Mb. Genome annotation predicted 34941 protein-coding genes with high-confidence. Using orthologous gene groupings, we constructed a phylogenetic tree comparing *G. glabra* with *G. inflata*, *G. uralensis*, and 10 other legume-related species. The analysis revealed that *G. glabra* shares the closest evolutionary relationship with *G. inflata*. Building on this, we then conducted a systematic genome-wide annotation, with a particular focus on candidate tailoring genes. Moreover, we performed a phylogenetic analysis to functionally annotate key reductases-PTR, VR and IFR.

**Figure 2.**
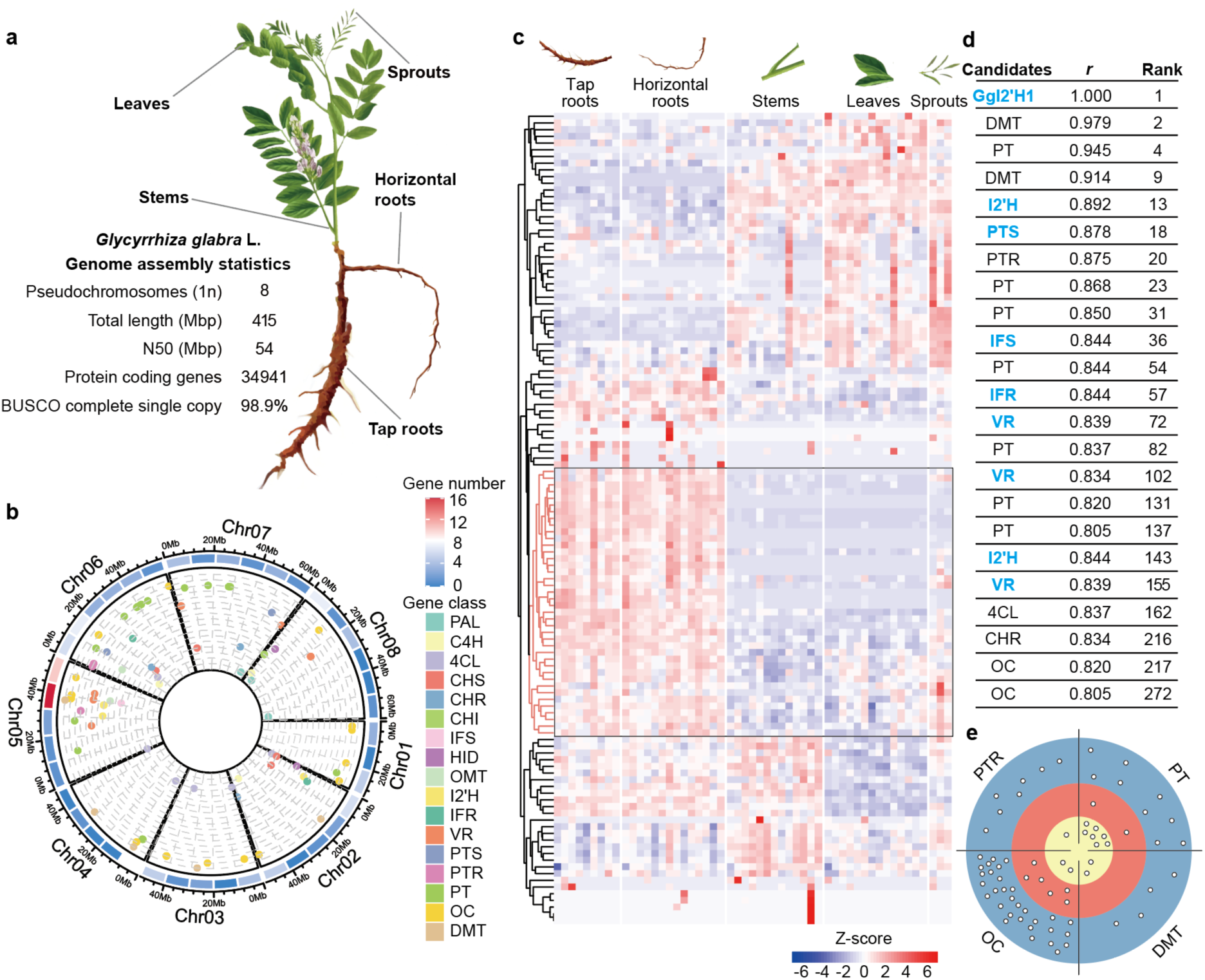
Genomic and transcriptomic analysis of *G. glabra* resources. a) *G. glabra* pseudochromosome genome assembly and gene annotation statistics. b) Genomic distribution of biosynthetic enzyme candidates for glabridin in *Glycyrrhiza glabra*. The outermost layer indicates the density of candidate genes across the genome (measured as number of genes per 1Mb). c) The transcriptional profiles of these genes across five tissues of *Glycyrrhiza glabra*: tap roots, horizontal roots, stems, leaves and sprouts (121 genes compared across 53 tissue samples). The expression levels are represented as Z-scores derived from log_2_-normalized TPM. A specific cluster enriched in root-expressed genes is highlighted in red. d) Co-expression analysis of the *G. glabra* transcriptome using *Gg*I2’H1 as a bait gene. The co-expressed genes were ranked based on their Pearson’s correlation coefficient (PCC) with *Gg*I2’H1. Genes shown in black are newly identified glabridin biosynthetic genes (this study). Homologs of the blue-labeled genes (I2’H, IFR, VR, and PTS) were previously identified (*20*). e) Identification of candidate genes for four tailoring enzymes involved in glabridin biosynthesis. The dots represent individual candidate genes. Blue circles indicate all annotated candidates identified through annotation; red circles represent candidates with high expression in root tissues; yellow circles denote genes that co-expressed with *Gg*I2’H1.

To investigate the transcriptional patterns of genes involved in glabridin biosynthesis, we conducted a large-scale transcriptomic study across major *Glycyrrhiza* resource in China. We sampled five *Glycyrrhiza* species, across seven locations, at four key seasonal growth stages, and from different plant organs, resulting in a total of 183 distinct samples. The clustering analysis revealed that the organ-specific transcriptome profiles displayed clear differences, grouping mainly into three clusters: 1) tap roots and horizontal roots, 2) stems and sprouts, 3) leaves. In contrast, the overall transcriptomic differences among species were relatively minor, with variations observed only in certain specific genes. Based on genome-wide alignments using *G. glabra* as the reference, genes that failed to match in *G. inflata* and *G. uralensis* were filtered out. Further clustering of expression patterns across organs and months highlighted key biosynthetic enzymes in *G. glabra*, including 7 PTRs, 18 PTs, 39 OCs, and 6 DMTs. Notably, these genes exhibited chromosomal clustering, primarily located on chromosomes 5 and 6 (Fig. 2b), and are primarily distributed in open and active chromatin regions—areas enriched in genes and transcriptionally active(*19*).

To further identify genes involved in glabridin synthesis, we performed hierarchical clustering to generate co-expressed patterns of transcripts across five organs (Fig. 2c). The clustering revealed a root-enriched gene expression cluster, including 1 PTR, 9 PTs, 8 OCs, and 2 DMTs. To refine candidate genes, we used previous reported key gene *Gg*I2’H1(*20*) as a bait for co-expression analysis. This analysis prioritized additional candidate enzymes, including 1 PTR, 7 PTs, 2 OCs, and 2 DMTs.

### Stepwise characterization of Glabridin biosynthetic key enzymes

To characterize the initial tailoring enzyme involved in the glabridin pathway, we first focused on pterocarpin reductase (PTR) that catalyzes the reductive cleavage of the dihydrofuran ring within potential core scaffolds, medicarpin (**8**) and demethylmedicarpin (**12**) (Fig. 3). We tested the one PTR candidate (*Gg*PTR1) identified through co-expression analysis. This enzyme convert medicarpin (**8**) and demethylmedicarpin (**12**) into vestitol (**9**) and demethylvestitol (**13**), with conversion rates of 90.6±3.6% and 52.7±0.9%, respectively (Fig. 3a). This confirms co-expression analysis as an effective strategy for prioritizing candidate enzymes. To identify more efficient PTRs, we further evaluated the remaining 6 PTR genes. Among them, the homolog *Gg*PTR5 exhibits detectable PTR activity but is less active than *Gg*PTR1. Notabley, *Gg*PTR2 with a different transcription pattern—highly expressing in the stems and leaves—also exhibits activity, with conversion rates of 58.8 ± 7.6% for compound (**8**) and 4.7 ± 2.5% for compound (**12**), respectively (Fig. 3b). In addition, although sharing 71% sequence similarity with *Gg*PTR1, *Gg*PTR2 can convert **25** to 4’-O-Methylpreglabridin (**10**), whereas *Gg*PTR1 shows no activity, indicating selective recognition of prenylated substrates by *Gg*PTR2 (Fig. 3a). To further explore the functional divergence, we modeled their structures in complex with medicarpin (**8**) and licoagrocarpin (**25**) (Fig. 3c). The electrostatic potential maps(*21*) indicated that both enzymes possess a positively charged catalytic pocket, attracting the electronegative phenolic hydroxyl groups of the substrates. Additionally, *Gg*PTR2 exhibited a similar binding orientation for both **8** and **25**, consistent with the binding of **8** in *Gg*PTR1. In contrast, compound **25** adopted a 180-degree flipped orientation in *Gg*PTR1, which may explain its lack of catalytic activity toward this substrate (Fig. 3c).

**Figure 3.**
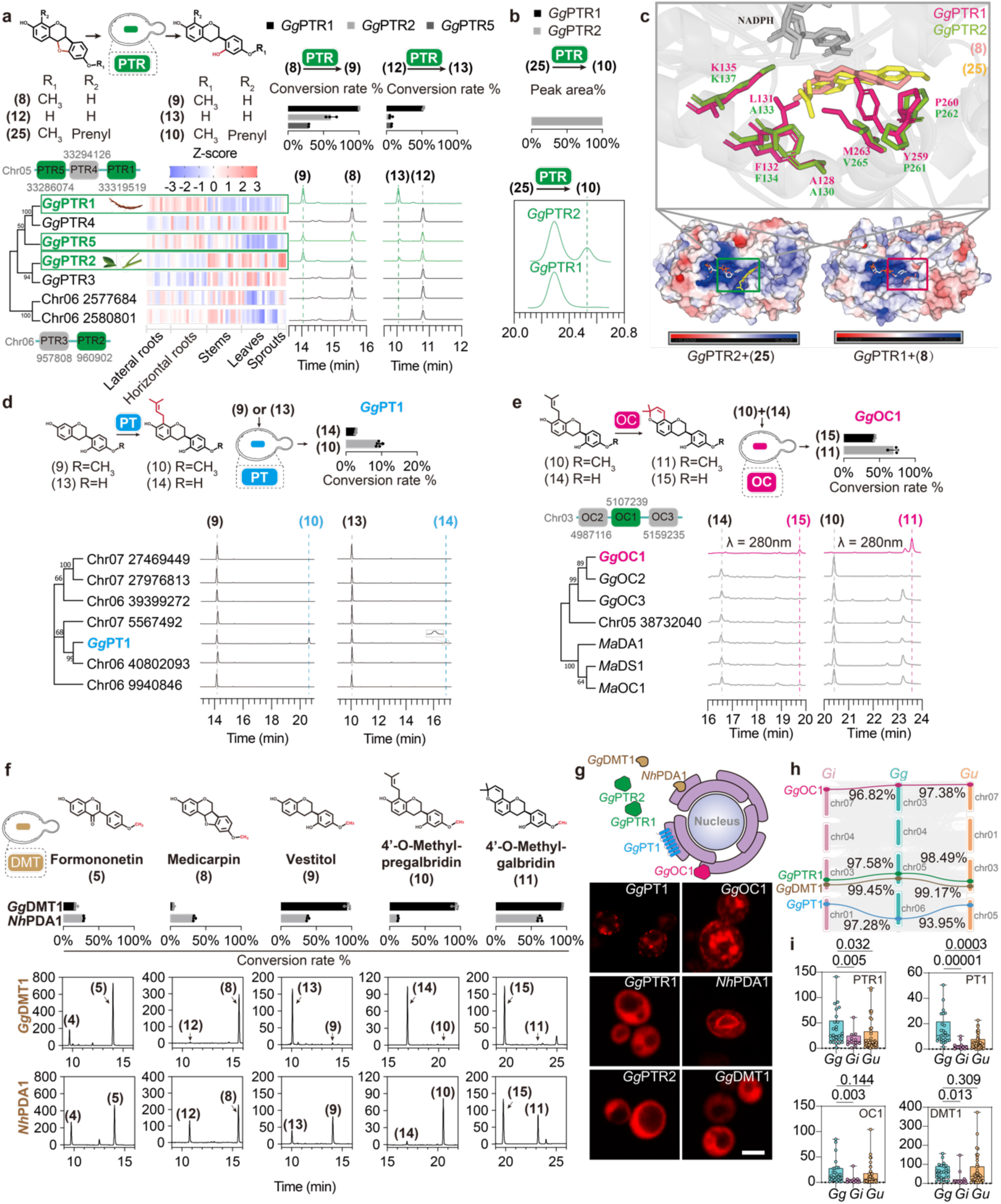
Characterization of the four tailoring enzymes. a) Catalytic activity characterizations of PTRs. The tissue-specific expression heatmap demonstrates the transcriptome data from tap root, horizontal root, stem, and leaf. Expression levels are log₂-transformed and normalized as Z-scores. b) Functional characterization of PTRs in catalyzing the conversion of prenylated pterocarpan substrate (**25**). c) Structural modeling of *Gg*PTR1 and *Gg*PTR2 in complex with substrates (**8** and **25**) using AlphaFold3(*37*). Boxed regions highlight major differences in the catalytic pocket. Electrostatic potential calculated using PyMOL APBS tools; Connolly surface shown within ±5 eV. Functional characterization and phylogenetic analysis of d) PTs and e) OCs. f) Substrate profiling of DMTs in catalyzing the demethylation of intermediate compounds. g) Yeast confocal microscopy showing subcellular location of mCherry-*Gg*PT1, *Gg*OC1-mCherry, *Gg*PTR1-mCherry, *Nh*PDA1-mCherry and *Gg*DMT1-mCherry. Scale bar, 5 μm. Images are representative of three independent experiments. Comparison of h) protein sequence homology and i) transcriptional expression levels of *Gg*PT1, *Gg*OC1, *Gg*PTR1, and *Gg*DMT1 across three *Glycyrrhiza* species (*G. glabra, n=*26; *G. inflata*, n=12; *G. uralensis,* n*=*39) in roots. Box plots in i.: lines are median, box limits are quartiles 1 and 3, whiskers are 1.5×interquartile range and points are outliers. The bar chart shows the molar conversion rates in a., d., e., and f.. Functional assays were performed by feeding 20 mg/L substrate in a. and 50 mg/L substrate in d., e., and f., and conversion rates were determined based on the relative peak areas of the corresponding products. Data represent the mean of at least three biological replicates.

Next, to identify prenyltransferases of *G. glabra*, we tested the seven PT candidates and found *Gg*PT1 exhibits detectable activity, catalyzing the conversion of compounds **9** and **13** into 4’-O-methylpreglabridin (**10**) and preglabridin (**14**) (Fig. 3d). Other PT genes, despite sharing a certain degree of phylogenetic similarity with *Gg*PT1, show no catalytic activity toward the tested substrates (Fig. 3d). Besides the enzymes from *G. glabra*, a reported *Pc*M4DT can catalyze the production of **10** from **25**(*22*).

After specific prenylation, intramolecular cyclization is required to form the characteristic cyclic ether structure(*23–26*). We then identified the two genes, *GgOC1* and *GgOC2*. Interestingly, the *GgOC1* exhibits an asymmetric co-expression pattern: the expression of *OC* is highly dependent on the transcriptional activity of the upstream genes(i.e. *GgI2’H1*), whereas the upstream genes show relatively weak association with *OC* expression. This strongly suggests that the *Gg*OC1 is most likely to participate in glabridin biosynthesis. Indeed, functional assays of the two OCs demonstrated that only *Gg*OC1 catalyzes the oxidative cyclization of 4’-O-methylpreglabridin (**10**) and preglabridin (**14**), yielding 4’-O-methylglabridin (**11**) and glabridin (**15**) (Fig. 3e). Further analysis revealed that *Gg*OC1 belongs to the BBE-type enzyme family and clusters with previously reported *Ma*OC1 in phylogenetic tree, which is responsible for the oxidative cyclization of prenylated chalcones(*27*).

Next, to find the DMTs in licorice species to realize the demethylation tailoring, we identified a gene ranked second, *Gg*DMT1 from the 2 candidates. It encodes a 2-oxoglutarate-dependent dioxygenase and reveals the ability to completely realize the demethylation of compound **11** (Fig. 3f). Interestingly, when checking the substrate profiling, we found *Gg*DMT1 shows high promiscuity and efficiently demethylates both vestitol (**9**) and 4’-O-methylpreglabridin (**10**). It also shows moderate activity toward formononetin (**5**), but displays very weak on medicarpin (**8**) (Fig. 3f). To further expand the enzyme toolbox for specific 4’-O-demethylation, we screened a panel of previously reported demethylases, including cytochrome P450s enzymes(*28–32*) from plant, fungal and human. Among these, only the fungal enzyme *Nh*PDA1 displayed demethylation activity, displaying promiscuity as well across toward multiple 4’-O-methylated substrates (**5**, **8**, **9**, **10**, **11**) (Fig. 3f). These observations are consistent with prior reports that demethylases often exhibit broad substrate specificity(*27, 32, 33*).

Given that *Gg*DMT1, originating from *G. glabra*, exhibits strong substrate promiscuity and high catalytic activity, it raises the question of why methylated intermediates still accumulate in this specie. Analysis of seasonal transcriptional patterns revealed an opposite trend between O-methyltransferase and demethylase expression, suggesting that precursor synthesis and final product accumulation may occur in temporally distinct stages. Furthermore, subcellular localization studies showed that *Gg*DMT1 is cytoplasmic, whereas *Gg*OC1 and *Gg*PT1 are more like a membrane-associated proteins (Fig. 3g). This spatial separation implies that the four tailoring enzymes may be compartmentalized—potentially within specific organelles— thereby facilitating intermediate segregation and stepwise regulation of the biosynthetic pathway(*34–36*).

Additionally, we found that throughout the pathway, methylation occurs first on the same hydroxyl group, followed by demethylation. This raises another question: whether such complexity is essential for pathway. When we examined alternative routes that bypass methylation reactions, we found that the I2’H1–IFR1–VR1 module is capable of catalyzing the conversion from intermediate **4** to **12**. However, its catalytic efficiency is extremely low. Molecular dynamics simulations revealed that (**17**) affects the reduction process by disrupting the hydrogen-bonding network between the cofactor NADPH and VR. These results suggest that methylation tailoring effectively blocks the non-methylated route and instead directs metabolic flux toward the more efficient methylated pathway.

Interestingly, when we analyzed the presence of these key enzymes in the other two *Glycyrrhiza* species, we found that both harbor homologous genes with >93% protein sequence identity, and their genomic loci are relatively conserved (Fig. 3h). We also analyzed the transcriptional differences of the four key enzymes across the three species and found that the transcription level of *PT* in *G. glabra* is significantly higher than in the other two species. *PTR*, *OC and DMT* also show significantly higher transcript levels in *G. glabra* compared to *G. inflata* (Fig. 3i). These findings help explain the significantly higher accumulation of glabridin in *G. glabra*, as observed in our metabolomic analyses (Fig. 1b). This may result from evolutionary changes in unknown regulatory networks that have activated the robust expression of glabridin biosynthetic genes in *G. glabra*, enabling the efficient production and accumulation of this bioactive compound.

### *De novo* biosynthesis of Glabridin in engineered yeast

To realize the biosynthesis of glabridin in yeast from the simple sugar glucose for the future industrilaztion, we designed the pathway with two modules, core scaffold module (glucose to **8**) and tailoring module (**8** to **15**) (Fig. 4a). We began by assembling the tailoring module using key enzymes we had characterized. Given that all involved enzymes-including PTR, PT, OC, and DMT-exhibit broad substrate promiscuity, we reconstructed the metabolic landscape from compound **8** to **15** and discovered a complex, interconnected tailoring network. This network supports at least two major tailoring routes: PT-PTR-OC and PTR-PT-OC. We then explored different enzyme combinations to further unravel this “tailoring maze”, aiming to realize glabridin production.

**Figure 4.**
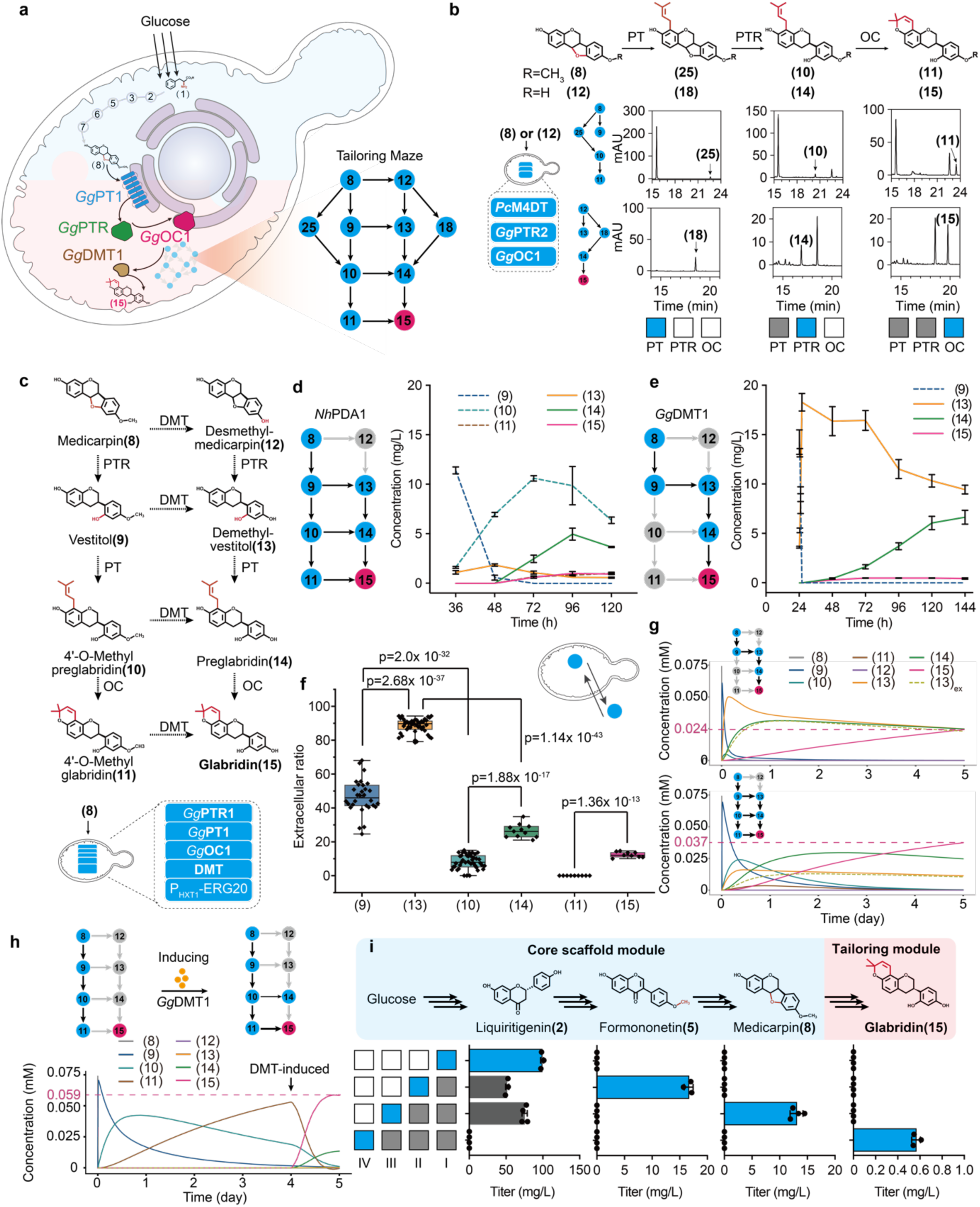
Glabridin biosynthesis module assembly and *de novo* production in yeast. a) Scheme of the glabridin biosynthetic pathway reconstructed in *S. cerevisiae*. b) Stepwise reconstitution of the glabridin biosynthetic pathway (PT–PTR–OC). 20 mg/L **8** and 10 mg/L **12** were added separately to engineered yeast strains with *Pc*M4DT-*Gg*PTR2-*Gg*OC1 combination. The squares on the bottom indicate different enzyme combinations: grey indicates enzymes present in the chassis, blue indicates newly introduced enzymes, and white indicates enzymes not introduced. c-e) Ladder-like biosynthetic network of glabridin (PTR– PT–OC–DMT). d-e) Metabolic dynamics of the ladder-like glabridin biosynthetic network incorporating with either d) *Nh*PDA1 or e) *Gg*DMT1. f) Extracellular distribution ratio of key intermediates in the ladder-like glabridin biosynthetic network. Box plots in f.: lines are median, box limits are quartiles 1 and 3, whiskers are 1.5×interquartile range and points are outliers. g) Metabolic simulation of multi-route biosynthetic network and single-route design. h) Metabolic simulation of induced pathway. i) *De novo* biosynthesis of glabridin in engineered *S. cerevisiae*.

We first tested the *Pc*M4DT-*Gg*PTR2-*Gg*OC1, which proceed as expected catalytic sequence: PT-PTR-OC, and continuous convert **8** to **11**, or **12** to **15** (Fig. 4b). Due to the promiscuity of *Gg*PTR2, side products such as **9** and **13** were also detected. In this design, we observed significant accumulation of intermediate **8**, indicating that the prenyltransferase reaction is inefficient—likely due to low the prenyl donor DMAPP. To address this, we dynamically regulated *ERG20* gene and overexpressed key rate-limiting enzymes (*tHMG1*, *IDI1*) for regulation of DMAPP supply.

In an alternative configuration PTR-PT-OC, we combined *Gg*PTR1-*Gg*PT1-*Gg*OC1, and when paired with a demethylation step, successfully reconstructed a ladder-like metabolic network. In this architecture, the shared enzymes-PTR, PT, and OC-act on different intermediates along two parallel metabolic routes, which are further interconnected by DMT, resulting in a flexible and interwoven biosynthetic network. Notably, the choice of demethylase significantly influenced the network’s behavior. Using *Nh*PDA1, a moderately active demethylase, the metabolic flux is distributed across multiple routes, forming a multi-route architecture in which various intermediates converge toward glabridin as the final product (Fig. 4d). This metabolism pattern is very close to that we detected in *G. glabra* (Fig. 1a-b), indicating *G. glabra* may contain a multi-route architecture as well. Moreover, we observed that flux preferentially flow toward compound **10**, while conversion to **13** remains low. After 48 hours, the demethylation of **10** became dominant, initiating glabridin production. This setup yields 0.9 mg/L of glabridin (Fig. 4d). However, compound **14** accumulates over time, indicating a bottleneck at oxidative cyclase (OC) step. In contrast, *Gg*DMT1, with its exceptionally high demethylation activity, drives the flux predominantly from compound **9** to **13**, effectively creating a streamlined, single-route pathway. In this mode, **10, 11** and **12** were not observed. When checking yield of glabridin, surprisingly, the simplified single-route design achieved much lower productivity than the complex, multi-route “ladder” network (Fig. 4), challenging the conventional view that flux distribution in multi-route systems reduces overall efficiency.

To investigate this phenomenon, we deeply analyzed metabolism dynamics of this network and observed intermediates secretion during fermentation (Fig. 4f). Specifically, prenylation reduces extracellular secretion, potentially due to increased hydrophobicity, while demethylation promotes secretion into the medium. These observations suggest that both methylation and prenylation act as self-retention strategies, helping to retain intermediates within the cell and minimizing loss through secretion. Among intermediates, **13** as a key metabolic node, demonstrates highest extracellular ratio, reaching 88.4±4.2% compared with the others. This extensive leakage explains the lower overall yield observed in the simplified single-route design. In contrast, despite with the strong efflux of **9** and **13**, multi-route network maintains higher level of robustness than the single one, suggesting the inherent redundancy and interconnectedness of the multi-route network provide robustness. To validate these observations, we built a metabolic flux model, which confirms the experimental trends and further support the network’s behavior (Fig. 4g). Interestingly, this model also indicates that shifting the demethylation step close to the end of the pathway could substantially improve glabridin yield, offering a promising strategy for optimizing *de novo* biosynthesis (Fig. 4h).

To achieve *de novo* biosynthesis of glabridin in *S. cerevisiae*, 14 enzymes were introduced to establish the core scaffold module (glucose to compound **8**), enabling the stepwise biosynthesis of liquiritigenin, formononetin and ultimately medicarpin. Specifically, seven genes-*Gg*PAL1, *Gg*C4H1, *Gg*4CL1, *Gg*CHS1, *Gg*CHR1, *Gg*CHI1 and *Sts*TAL-were incorporated to drive liquiritigenin biosynthesis from glucose. Liquiritigenin was subsequently converted to formononetin via three additional enzymes: *Gg*IFS1, *Gg*HID1, and *Gg*OMT1. To complete the formation of the core scaffold medicarpin, four more enzymes-*Gg*I2’H1, *Gg*IFR1, *Gg*VR1, and *Gg*PTS1-were expressed in yeast. Following establishment of the core pathway, the tailoring module (*Gg*PTR1-*Gg*PT1-*Gg*OC1-*Gg*DMT1) was introduced, with *Gg*DMT1 controlled by the cyanamide-inducible DDI2 promoter. In addition, to relieve limitations in the *de novo* biosynthesis of glabridin, the rate-limiting enzymes were overexpressed and NADPH supply were enhanced. Finally, this fully engineered yeast strain enabled *de novo* production of glabridin in shake-flask fermentation, reaching a final titer of 0.5 mg/L (Fig. 4i).

## Conclusion

In this study, we systematically mapped the maze-like biosynthetic network of glabridin and successfully reconstructed it in *S. cerevisiae*, achieving *de novo* production from glucose. By identifying and assembling key tailoring enzymes, we revealed a ladder-like network architecture mediated by a set of promiscuous enzymes that enhances pathway robustness through multiple interconnected routes. Unlike classical linear metabolic pathways, the multi-route design tolerates metabolic flux fluctuations(*38, 39*), contributing to improved overall productivity. Our study also provides new insights into the evolutionary adaptation of biosynthetic pathways in *Glycyrrhiza* species. The observed metabolic promiscuity and redundancy may represent an evolutionary strategy to maximize ecological adaptability while minimizing genetic burden, by utilizing a limited number of gene copies to ensure metabolic resilience and flexibility across diverse environmental conditions(*40, 41*).

While we have successfully reconstructed the major biosynthetic routes, several challenges remain for further pathway optimization. Notably, the oxidative cyclization step catalyzed by *Gg*OC1 exhibited low catalytic efficiency, emerging as the rate-limiting step for glabridin biosynthesis. Moreover, although we established two major biosynthetic routes, certain branches of the network (Fig. 1) remain incomplete due to missing or unidentified enzymes. Future efforts to discover and integrate these enzymes could fully realize the potential of the glabridin biosynthetic network, enabling more efficient and versatile production of glabridin and structurally related derivatives.

Overall, our work not only provides a blueprint for glabridin biosynthesis but also highlights a generalizable principle: maze-like, multi-route architectures can endow microbial cell factories with enhanced metabolic resilience, offering a new strategy for the synthesis of complex plant natural products. In addition, the development of such scalable and sustainable microbial production systems offers a highly promising alternative to direct plant extraction, which is essential for the wildlife conservation of rare plant populations, such as *Glycyrrhiza glabra*.

## Materials and Methods

### Chemical standards

All chemical standards used in this study were obtained either from commercial suppliers or through custom synthesis. Preglabridin was obtained from TargetMol (Boston, MA, USA). 4’-O-methylglabridin and 4’-O-methylpreglabridin were custom-synthesized by Acme Biopharma Co., Ltd. (Shanghai, China). All other chemicals were purchased from YuanYe Biotechnology Co., Ltd. (Shanghai, China).

### Retrobiosynthesis analysis

The bio-retrosynthesis algorithm was employed to predict the biosynthetic pathway of glabridin. First, the molecular fingerprint similarity method (FingerprintSimilarity) from the RDKit package in Python was utilized to identify structural analogs of glabridin among metabolites in the KEGG database (https://www.genome.jp/kegg/pathway.html). Second, the acyclic path search method (shortest_simple_paths function) from NetworkX package in Python was applied to explore potential biosynthetic routes for these structural analogs. The pathway origins were set as both the identified glabridin analogs and 2,808 endogenous metabolites from the *S. cerevisiae* genome-scale metabolic model YEAST8(*42*). Subsequently, RetroRules(*43*) (containing 65,688 reaction rules with diameters = 4 and 6) was implemented for retrosynthetic pathway prediction from glabridin to its structural analogs identified in the previous step. To address combinatorial explosion challenges, Monte Carlo Tree Search (MCTS) was employed to establish a retrosynthetic route from glabridin to its structural analog medicarpin. This foundation was further expanded through human-computer interaction single-step retrosynthetic analysis to develop a comprehensive retrosynthetic network.

### Genome sequencing, assembly and annotation

Samples of *Glycyrrhiza glabra* were collected from the Alar region of Xinjiang, China. A PacBio sequencing library was constructed and sequenced using six SMRT cells on the PacBio Sequel platform (Pacific Biosciences) by BGI Genomics Co., Ltd. The third-generation sequencing data were first corrected using the Canu(*44*) (v2.2), followed by preliminary assembly with the Smartdenovo(*45*) software (v1.0.0,) to obtain initial contig sequences. To further improve assembly quality, the initial contigs were sequentially polished using Racon(*46*) (v1.4.3) for third-generation data correction, followed by Pilon(*47*) (v1.24) for error correction with second-generation sequencing data, yielding the final preliminary assembly. To remove redundant regions, the genome assembly was processed using the Purge haplotigs(*48*) (v1.1.2) tool. Genome completeness was assessed using BUSCO(*49–51*) (v5.8.3). A Hi-C sample was sequenced for high-quality data. By integrating Hi-C linkage signals, the contig sequences were oriented, clustered, and anchored to the chromosomal level using Juicer(*52*) (v1.6) and 3d-dna (v201008) (*53*). Illumina and PacBio sequencing reads were mapped on chromosome level genome assembly with Bowtie2 (v2.5.4) and minimap2 (v2.28), respectively, where default parameters were adopted (*54, 55*). Then sequencing coverage was calculated by ‘depth’ function of Samtools (v1.18) in ‘-aa’ mode and displayed in a Circos plot with TBtools (v2.007) (*56, 57*). Phylogeny relationship of *G. glabra*, *G. inflata*, *G. uralensis* and thirteen neighbor species in Leguminosae were constructed according to profiles of shared and unique orthology genes with OrthoFinder (v2.5.4) by default parameters(*58*).

To annotate the SDR family enzymes identified from licorice, enzymes from the SDR406A family were retrieved using the Uniprot database (https://www.uniprot.org). The BLAST tool (https://blast.ncbi.nlm.nih.gov/Blast.cgi) was used to search for homologous sequences of the target enzymes. From the identified candidates, enzymes with known functions were selected, and a multiple sequence alignment was performed using MUSCLE(*59*) (version 5.1). Phylogenetic tree construction was carried out using IQ-TREE(*60*)(version 2.1.4-beta), applying the Maximum Likelihood (ML) method. The inference of the phylogenetic tree was based on the JTT (Jones-Taylor-Thornton) amino acid substitution model, with 1000 ultrafast bootstrap(*61*) resampling to assess branch support and an additional 1000 approximate likelihood ratio tests to further evaluate branch reliability. The final tree was visualized using online iTOL (https://itol.embl.de), with clusters containing only branches that include the enzymes to be annotated retained. The branch lengths of the tree represent evolutionary distances.

### Total RNA isolation, RNA-Seq and gene expression quantification

Based on the geographical distribution of wild licorice resources, sampling sites were selected in Turpan, Urumqi, Emin, Tumxuk, Aksu in Xinjiang, and Longdong in Gansu. From the same individual plant at each location, distinct samples of tap roots, horizontal roots, stems, leaves and sprouts were collected. To facilitate subsequent metabolite analysis and RNA extraction for transcriptome profiling, the collected tissues were immediately flash-frozen in liquid nitrogen. Aliquots intended for metabolomic analysis were stored on dry ice, while aliquots for transcriptomic analysis were preserved in an RNA stabilization solution. Licorice was chopped and ground by freezing with liquid nitrogen. Total RNA was extracted by using the TranZol™ kit (TransGen Biotech, China) according to the manufacturer’s instructions, and was subsequently used to synthesize first-strand complementary DNA (cDNA) with the TransScript® One-Step gDNA Removal and cDNA Synthesis SuperMix (TransGen Biotech, China). Transcriptome sequencing of various *G. glabra* tissues was performed by BGI Genomics Co., Ltd. (Shenzhen, China). The quality of RNA was assessed by the Agilent 2100 Bioanalyzer, and mRNA was purified using oligomagnetic beads. The reverse-transcribed cDNA library was sequenced using BGI’s BGISEQ500RS platform (BGI Genomics Co., Ltd). The raw data and measured parameters of the transcriptome have been stored in Jingweikaiwu database (kw.ibc.tsinghua.edu.cn). For transcriptome analysis, RNA-seq reads were aligned to the genome sequence using Hisat2 (v2.2.1)(*62, 63*). Subsequent gene prediction was performed using GeMoMa(*64, 65*), leveraging genomic and transcriptomic data from a closely related species. The gene set was evaluated for completeness using BUSCO. For transcriptome quantification, transcript abundance was quantified using Stringtie(*66*).

### Co-expression analysis of candidate genes

A heatmap was generated to visualize the transcriptional profiles of candidate genes across five tissues of *Glycyrrhiza glabra*—tap roots, horizontal roots, stems, leaves and sprouts. The heatmap was constructed using the R package pheatmap (v1.0.12), with row-wise scaling. Expression levels were shown as Z-scores calculated from log_2_-normalized TPM values. Pearson correlation analysis was performed using *Gg*I2’H1 as the bait gene, based on TPM values across 53 *G. glabra* transcriptomes covering the five tissues.

### Plasmid construction

Codon-optimized synthetic genes were synthesized by Tsingke (Beijing, China). Two classes of plasmids were constructed: functional assay plasmids and pathway integration plasmids. For functional assays, candidate genes were cloned between a galactose-inducible GAL2 promoter and the ECM10 terminator, flanked by 500–800 bp genomic homology arms to enable targeted integration into the yeast genome. All vector components except for the gene insert were synthesized in advance. A pair of inward-facing BsaI restriction sites was placed between the promoter and terminator to allow seamless gene insertion via Golden Gate assembly. Gene fragments (either PCR-amplified or synthesized) were digested with BsaI and ligated using T4 DNA ligase (both from New England Biolabs). Pathway integration plasmids were constructed using the same BsaI-based cloning strategy. In these plasmids, expression cassettes consisted of GAL2 or GAL1 promoters from various species, and native yeast terminators. Each expression cassette was flanked by 200 bp synthetic orthogonal sequences that were manually designed to serve as overlap regions, enabling one-step assembly of multi-gene constructs.

### Yeast Strain construction

Yeast genomic modifications were performed using the CRISPR-based method(*67*). DNA fragments intended for genomic integration were PCR-amplified using Q5 DNA polymerase (NEB) and co-transformed with a CRISPR-Cas9 plasmid targeting the desired locus. For yeast transformations, an overnight culture of the parental strain was inoculated into 4 mL of 2×YPD medium in a 24-deep well plate at an initial OD_600_ of 0.2 and incubated at 30 °C with shaking at 400 rpm until OD_600_ reached 1.0. Cells corresponding to 1 OD unit were harvested by centrifugation at 3,000 × g for 2 min and washed with an equal volume of sterile water. The cell pellet was then resuspended in a transformation mixture containing 1,000 ng of DNA fragment, 500 ng of CRISPR-Cas9 plasmid, and 260 μL of 50% PEG3350, 36 μL of 1 M LiOAc, and 10 μL of denatured salmon sperm ssDNA(*68*). The mixture was incubated at 42 °C for 30 min. Following centrifugation at 3,000 × g for 2 min, the cell pellet was resuspended in 600 μL of YPD and recovered at 30 °C for 90 min. Cells were then pelleted again, resuspended in 100 μL of sterile water, and plated onto selective agar plates. Correct integration events were confirmed by colony PCR. Verified clones were used for subsequent engineering steps after curing the CRISPR-Cas9 plasmid. Yeast culture media were obtained from Coolaber (Beijing, China).

### Strains and culture conditions

Yeast colonies were initially inoculated in triplicate into 2 mL YPD and grown overnight to saturation at 30 °C and 400 rpm, and then 100 μL transferred into 4 mL deep-well 24-well plates sealed with Breathable Sealing Film (Biotss). Cultures were incubated for 120 h at 30 °C, 400 rpm in a high-speed orbital shaker (ZSZY-88BH; Shanghai Zhichu Instrument Co., Ltd.). After 24 h of incubation, galactose was added to a final concentration of 20 g/L to induce gene expression, along with substrate addition when required.

### Metabolomic profiling of licorice

Dried roots, stems, and leaves of *Glycyrrhiza glabra* were ground into coarse powder under liquid nitrogen and further disrupted using 0.5 mm and 3 mm zirconia beads in a cryogenic bead mill (Guangzhou Luka Sequencing Instrument Co., Ltd.) at 75 Hz for five cycles (each cycle: 60 s grinding, 30 s pause) to obtain fine plant powder. Accurately weighed samples were extracted three times with 25 mL of methanol and 25 mL of ethyl acetate, respectively, in an ultrasonic bath at 25 °C for 30 min. The combined extracts were concentrated to dryness, re-dissolved in methanol, and filtered through a 0.22 μm membrane.

Metabolite profiling was performed using a Thermo Scientific Ultimate 3000 UHPLC system coupled to a Q Exactive Plus Orbitrap high-resolution mass spectrometer (Thermo Fisher Scientific, USA). Chromatographic separation was carried out on a Waters ACQUITY UPLC HSS T3 C18 column (2.1 × 100 mm, 1.8 μm; Waters Corporation, USA), with 0.1% (v/v) formic acid in water as mobile phase A and 0.1% (v/v) formic acid in acetonitrile as mobile phase B. The column temperature was maintained at 35 °C, the flow rate was set to 0.2 mL/min, and the injection volume was 5 μL. The elution was performed using the following gradient: 0.00–10.00 min, 100% B; 10.00–20.00 min, 100–70% B; 20.00–25.00 min, 70–60% B; 25.00–30.00 min, 60–50% B; 30.00–40.00 min, 50–30% B; 40.00–45.00 min, 30–0% B; 45.00–60.00 min, 0% B; 60.00–60.10 min, 0–100% B; 60.10–70.00 min, 100% B. Mass spectrometry was conducted in both positive and negative ionization modes using a heated electrospray ionization (HESI) source. The sheath gas flow rate was set to 40 arbitrary units, auxiliary gas to 15 units, capillary temperature to 320 °C, and auxiliary gas heater temperature to 350 °C. The spray voltages were +3.2 kV (positive mode) and –3.0 kV (negative mode). The resolution was set to 70,000 for MS and 17,500 for MS/MS. Data were acquired in full scan mode over an m/z range of 100–1500. Compound identification was performed using Compound Discoverer 3.3 software, with reference to mzCloud and mzVault databases. The peak Area of each detected compound was normalized to the corresponding sample weight and used for metabolomic heatmap visualization.

### Analysis of metabolite production

Yeast cultures were centrifuged at 3,000 g for 5 min to separate the cell pellets and supernatants. Cell pellets were resuspended in saturated saline and mixed with 0.5 mm zirconia beads. Cell disruption was performed using a cryogenic bead mill (Guangzhou Luka Sequencing Instrument Co., Ltd.). Ethyl acetate was added to both the disrupted cell suspension and the culture supernatant for metabolite extraction. The organic phase was collected and evaporated to dryness under vacuum using a concentrator (Concentrator plus; Eppendorf). Residues were re-dissolved in methanol and filtered through a Nylon 66 membrane (Bioexploration, Guangdong, China) to obtain samples for subsequent HPLC analysis. Metabolites were analyzed using an Agilent 1260 Infinity Binary HPLC system. Chromatographic separation was performed on an InfinityLab Poroshell 120 EC-C18 column (3.0 × 100 mm, 2.7 μm; Agilent Technologies) with 0.1% (v/v) formic acid in water as mobile phase A and acetonitrile as mobile phase B. The flow rate was set to 0.5 mL/min, with a column temperature of 40 °C and an injection volume of 5 μL. The elution was carried out using the following gradient: 0.00–5.00 min, 10–20% B; 5.00–20.00 min, 20–60% B; 20.00–32.00 min, 60–85% B; 32.00–34.00 min, 85–100% B; 34.00–36.00 min, 100% B; 36.00–38.00 min, 100–10% B; 38.00– 40.00 min, 10% B. LC–MS analysis was performed following the same procedure described in the “Metabolomic profiling of licorice” section. The production of metabolite was purified and characterized by liquid Chromatography-Mass Spectrometry (LC-MS) and structural determination by NMR.

### Protein structure modeling and docking

Protein structures and ligand docking were predicted using the local version of AlphaFold 3(*69*). The SMILES notation of the ligands was obtained from the PubChem database (https://pubchem.ncbi.nlm.nih.gov). PyMOL (version 2.5.7, http://www.pymol.org) molecular visualization software was used for further analysis of the ligand-receptor complexes. The APBS Electrostatics feature in PyMOL was applied to calculate the electrostatic potential maps of the proteins, utilizing the pdb2pqr method with a grid spacing of 0.50 Å. The predicted isoprenoid-binding pockets were determined by comparing protein structures and sequence similarities.

### Molecular dynamics simulation

The force field parameters of NADPH were retrieved from previous reports(*70*). The geometry of the ligand was optimized using the ORCA 6.0(*71, 72*) with the M06-2X method(*73*) combined with the def2-TZVP(*74, 75*) basis set and empirical dispersion correction GD3(*76*). Molecular force field parameters for the ligand were generated by the sobtop tool based on the General Amber Force Field (GAFF2)(*77, 78*), with atomic charges fitted using the RESP method(*79*). Protein topology was described using the Amber ff14SB all-atom force field(*80*). Molecular dynamics (MD) simulations were conducted using the GROMACS 2023 software package(*81*). The protein-ligand complex was placed in a periodic boundary condition box filled with the TIP3P water model(*82*), with the system solvated and neutralized by adding 0.1 M NaCl. The simulation system underwent sequential energy minimization, followed by equilibration under the NVT ensemble at 303.15 K and the NPT ensemble. Each complex adopts three independent 200 ns simulations with a time step of 2 fs, and trajectories were recorded every 100 ps. System temperature was maintained at 303.15 K using the V-rescale thermostat, while pressure was controlled at 1 bar via the C-rescale method. Non-bonded van der Waals interactions were computed using a switching function with a cutoff of 1.0 nm, and long-range electrostatic interactions were calculated with the particle mesh Ewald (PME) method(*83*), also using a cutoff of 1.0 nm. The distances between atoms were calculated by grabbing atoms from the index file with the “gmx distance” command to determine distance variations. Hydrogen bond occurrences were quantified using the “gmx hbond” command. Simulation results were visually analyzed using PyMOL software (version 2.6.2).

### Fluorescence image and analysis

Individual colonies of yeast strains transformed with plasmids encoding biosynthetic enzymes fused to fluorescent protein reporters were inoculated into 4 mL YPD medium and grown overnight (∼12 h) at 30 °C with shaking at 400 rpm. Overnight cultures were then diluted 1:4 into fresh YPD medium and incubated for an additional 6–8 h at 30 °C, 400 rpm to allow proper folding of slow-maturing fluorescent proteins. Cells were harvested by centrifugation and resuspended in 100 μL of sterile PBS for imaging. For microscopy, 5–10 μL of cell suspension was spotted onto a glass microscope slide and covered with a glass coverslip (Shanghai Titan Scientific Co., Ltd.). Imaging was performed using a Nikon AXR multi-SIM multimodal super-resolution confocal microscope (Nikon Instruments Inc., Japan) equipped with a 100×oil immersion objective. Fluorescence excitation was carried out using 488 nm for the green channel (GFP) and 561 nm for the red channel (mCherry). Images were acquired using NIS-Elements software and processed using ImageJ (v1.54f).

### Ladder network dynamics measurement

Yeast colonies were initially inoculated in triplicate into 4 mL YPD medium and grown overnight at 30 °C with shaking at 400 rpm until saturation. Subsequently, 1 mL of each culture was transferred into 20 mL fresh YPD medium in 100-mL shake flasks. Cultures were incubated for 120 h at 30 °C and 250 rpm in a shaking incubator. After 24 h of incubation, galactose was added to a final concentration of 20 g/L to induce gene expression, along with substrate addition when applicable. Samples were collected every 24 h and processed using the same method described above.

### Model Framework and Simplifications

A computational modeling approach was employed to describe metabolic networks formed by multiple promiscuous enzymes and their substrate compounds. To address the inherent complexity of cellular metabolic environments, essential simplifications were implemented throughout the simulations. Cellular compartmentalization and compound diffusion rates were neglected in the model formulation.

The Michaelis-Menten equation was adapted to simulate catalytic processes, accounting for competitive interactions between: (1) different substrates for shared enzyme active sites, and (2) multiple enzymes competing for identical substrates. This framework yielded:

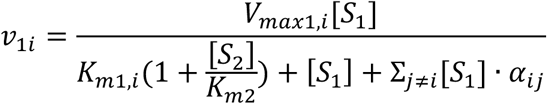

Km values for Michaelis-Menten equation were predicted through reported model (*84*). Computational analysis revealed Km ≫ [S], where substrate concentrations ([S]) were significantly lower than corresponding Km values. This permitted simplification of the Michaelis-Menten equation to:

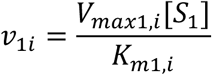

To align with laboratory conditions for microbial cell factories, the model assumed constant cellular biomass and enzyme concentrations. Enzyme concentrations were fixed based on biomass measurements at 24 hr post-inoculation, coinciding with substrate supplementation timing in experimental protocols.

Additionally, concerning the critical parameter of compound transport across cellular membranes (influx/efflux) that significantly impacts this reaction network, we postulate that all compounds achieve rapid equilibrium through passive transport mechanisms, represented by:

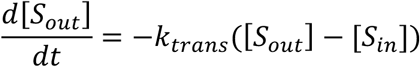

where transport rate constants (*k_trans_*) were experimentally determined through separate permeability assays.

The resulting system of ordinary differential equations was numerically solved using LSODA (Low-order SObolov differential equation solver), solve_ivp function in Python (v3.8.5, scipy 1.10.1).

### Quantification and statistical analysis

Data were processed by Excel, GraphPad Prism and R. The significant comparisons of two different groups are indicated in the graphs statistical analysis was performed using two-tailed unpaired Student’s t-test. P values were given in the figures. The graphs represented means ± SD unless otherwise indicated, as described in the figure legends.

## Acknowledgements

This work was supported by the Natural Science Foundation of China (22138006, 22278240, 22478032, 22478031).

## Author contributions

C.L., H.J., L.Q., and B.L. conceived the project. Z.Z. contributed to experimental design, planning, and execution. WQ.L. and W.L. performed genome assembly and analysis. B.L., W.S., S.R., and L.Z. collected Glycyrrhiza samples. L.Q. and Z.Z. analyzed the transcriptome data. FZ. M. developed the theoretical model. Q.K. and Z.Z. performed metabolomic analysis. Y.J. carried out enzyme structural modeling. WQ.L. and Y.J. performed enzyme kinetic simulations. All authors contributed to data analysis and interpretation of results. H.J. and Z.Z. drafted the manuscript. H.J., L.Q., B.L., and C.L. revised the manuscript and supervised the study.

## Competing interests

The authors declare no competing interests.

## Notes

### Competing Interest Statement

The authors have declared no competing interest.

### Summary of Updates

We have revised the some typo errors in the main text and in the fig 1.

